# PLUG (Pruning of Local Unrealistic Geometries) removes restrictions on biophysical modeling for protein design

**DOI:** 10.1101/368522

**Authors:** Mark A. Hallen

## Abstract

Protein design algorithms must search an enormous conformational space to identify favorable conformations. As a result, those that perform this search with guarantees of accuracy generally start with a conformational pruning step, such as dead-end elimination (DEE). However, the mathematical assumptions of DEE-based pruning algorithms have up to now severely restricted the biophysical model that can feasibly be used in protein design. To lift these restrictions, I propose to <u>p</u>rune <u>l</u>ocal <u>u</u>nrealistic <u>g</u>eometries (PLUG) using a linear programming-based method. PLUG’s biophysical model consists only of well-known lower bounds on interatomic distances. PLUG is intended as pre-processing for energy-based protein design calculations, whose biophysical model need not support DEE pruning. Based on 96 test cases, PLUG is at least as effective at pruning as DEE for larger protein designs—the type that most require pruning. When combined with the LUTE protein design algorithm, PLUG greatly facilitates designs that account for continuous entropy, large multistate designs with continuous flexibility, and designs with extensive continuous backbone flexibility and advanced non-pairwise energy functions. Many of these designs are tractable only with PLUG, either for empirical reasons (LUTE’s machine learning step achieves an accurate fit only after PLUG pruning), or for theoretical reasons (many energy functions are fundamentally incompatible with DEE).

## Introduction

Computational protein design is the identification of protein sequences that optimize a desired function, typically involving binding^6,9,31,37^. Specifically, given the 3-D structure of a protein, protein design algorithms can identify mutations in the sequence of the protein that will minimize its energy—either the energy of the protein itself^5,6,10,15,27^, or a derived quantity such as a binding energy of the protein to a ligand^7,13,25,30^. Predictions of sequences’ energies are based on a *biophysical model*. This model includes an energy function mapping molecular geometry to energy, as well as a set of conformational degrees of freedom that are considered flexible, and bounds on how much each degree of freedom can move.

Efficient protein design algorithms must search an enormous space of sequences and conformations, which grows exponentially as mutable residue positions are added to the design. Approximations are thus commonly made to speed up the search, at a significant cost in accuracy. One common assumption is that each residue can only access a limited, discrete set of conformations known as *ideal rotamers*^5,23^, which correspond to observed modal values in the space of sidechain dihedral angles. Another is that the energy function is a sum of terms depending on at most two residues’ conformations—that is, it is *pairwise*. The most efficient protein design algorithms rely on these assumptions to calculate an *energy matrix* that stores precomputed energetic interactions between pairs of rotamers at different residues. This precomputation dramatically speeds up protein design calculations. An energy matrix can be used to efficiently perform simulated annealing^4,27,48^, which can yield an answer quickly with no guarantees of accuracy and with error that empirically increases for larger designs^44^, or as input to a search algorithm with provable guarantees of accuracy. Many algorithms with such guarantees are available, including DEE/A*^5,10,13,17,22,29,36^, integer linear programming^26,40^, branch-^24^ and tree^50^ decomposition-based methods, and weighted constraint satisfaction algorithms^40,46,47^.

Although simulated annealing and DEE/A* have both been extended to account for continuous flexibility, this comes at a significant computational cost. Simulated annealing becomes very inefficient when the energy matrix is not available. Hence continuous flexibility, if accounted for at all, is used only in a small number of rounds of sampling at the end of the calculation^48^, which may prevent the identification of favorable conformations significantly different from the best conformation found in discrete search.

DEE/A* has been extended to account for continuous flexibility while maintaining guarantees of accuracy^10,13,20,21,38,39^, by using lower bounds on pairwise interaction energy that are calculated by minimizing pairwise interaction energies for particular rotamer pairs in isolation. In particular, while DEE determines if a rotamer can be pruned by computing a difference between energies based on a discrete energy matrix and checking if it is greater than zero^5^, the iMinDEE algorithm computes the same energy difference using the pairwise energy matrix and checks if it is greater than an energy buffer called the *pruning interval*^10^. iMinDEE is proven to be accurate if the pruning interval is large enough. It must be at least the difference between the globally optimal energy computed using the pairwise minimized energy matrix, and the true globally optimal energy (calculated using minimization of fully defined conformations rather than just pairs). Thus the pruning interval is in some sense an upper bound on how much the pairwise minimized energy can deviate from the true minimized energy, for purposes of pruning.

The machine learning-based LUTE algorithm^21^ is currently the most efficient way to perform protein design with continuous flexibility. It learns an energy matrix that closely approximates the continuously minimized energies. LUTE can be applied to multiple states of a protein system—for example bound and unbound states, possibly with multiple binding partners—to perform *multistate designs* that optimize for properties like binding affinity and specificity, using the COMETS algorithm^18^. However, the presence of high-energy conformations in a conformational space can cause the number of parameters required for a low-residual LUTE fit to be prohibitively large. Thus, a pruning method like iMinDEE must be performed before the fitting of the LUTE matrix. This limits the applicability of LUTE, and indeed of continuously flexible protein design in general, to designs that are either suitable for running iMinDEE pruning (requiring pairwise energy functions and certain other conditions) or have small enough sequence and conformational spaces to allow efficient and accurate LUTE fits with only very obvious steric clashes being pruned^21^. Thus, a more general pruning method would help for such designs.

PLUG provides this more general pruning method. Rather than relying on an energy matrix or indeed any biophysical assumptions about energy, it merely generates geometric constraints representing steric clashes (pairs of atoms coming unrealistically close together). The conditions that cause steric clashes have actually been characterized more precisely than protein energetics in general, and are routinely used in visualizing protein structures and detecting problems in crystallographic structure determination, e.g. using the Probe software^49^ and the MolProbity server^2^. PLUG eliminates clashing portions of conformational space from consideration in designs, enabling more efficient and accurate application of LUTE and other protein design algorithms.

PLUG is implemented in the osprey^11,13,14,35^ open-source protein design package. osprey has yielded many designs that performed well experimentally—*in vitro*^1,8,12,16,38,43,45^ and *in vivo*^8,16,38,43^ as well as in non-human primates^43^—and contains a wide array of flexibility modeling options and provably accurate design algorithms^11,14^. These features will allow PLUG to be used for many different types of designs.

By presenting PLUG, this paper makes the following contributions:

1. A method for pruning portions of protein (or other polymer) conformational space based on geometry and a model of steric clashes.
2. A method to apply this pruning method in protein designs using the LUTE algorithm, greatly facilitating its use with extensive backbone flexibility, non-pairwise energy functions, and continuous sidechain entropy and in multistate designs. Many of these designs would not be practical without PLUG, as shown empirically here.
3. Empirical results on 96 design test cases using 36 protein structures, demonstrating that PLUG’s pruning power is comparable to the previous state of the art, while it allows the use of a wider range of biophysical models in protein design.
4. An implementation of PLUG in the open-source osprey^11,13,14,35^ protein design software package, available at www.github.com/markhallen369/OSPREY_refactor

## Materials and Methods

### Background: Voxel representation of conformational space

Consider a polymer (for example, a protein) consisting of a chain of *residues*. Chemically, the polymer is defined by its *sequence*, which defines each residue as a particular chemical species (a particular amino-acid type, in the case of a protein). Designing the polymer requires searching over a sequence space, represented as a set of possible chemical species for each residue. For example, a simple sequence space for a protein might consist of all sequences in which a protein has either valine or alanine at residue position 1, either glutamate or aspartate at residue 2, and the wild-type amino-acid type at all other residues. The objective function for the search over sequence space can vary from design to design. One may wish to simply minimize the energy of the protein (to optimize stability), or optimize binding to a particular partner, or optimize specific binding to one partner but not another. Each of these types of designs can be approached using biophysical modeling, which requires a representation of the polymer’s conformational space: the set of 3-D geometries that each sequence can exhibit^18^. The conformational flexibility of the polymer will be modeled using *n* internal coordinates x = {*x_i_* | *i* ∊ {1,…, *n*}}. These internal coordinates can be sidechain dihedral angles or backbone perturbations^22^, or even backbone internal coordinates generated using the CATS algorithm^19^ that allow the protein backbone to move locally in all locally feasible directions. The conformational space of the system is then defined as the the union of *voxels*^10,13,22^. Each voxel *v* is defined by a sequence and the inequality constraints

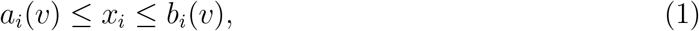

for *i* ∊ {1,…, *n*}, where *a_i_*(*v*) and *b_i_*(*v*) are voxel-specific bounds defined per the modeling assumptions. If *a_i_*(*v*) < *b_i_*(*v*), the coordinate *x_i_* has *continuous flexibility* in *v*.

The role of PLUG will be to identify voxels with continuous flexibility that are not biophysically feasible, and to exclude them from the conformational space. This simplifies the problem and enables the subsequent use of algorithms (e.g., LUTE^21^) that can search over a very large space of voxels as long as those voxels are “well-behaved” energetically. For example, when a conformation contains non-bonded atoms that are very close to each other, the energy is extremely sensitive to small conformational changes, leading to an ill-conditioned search problem that is difficult for LUTE to solve. But conformations with such atoms are not biophysically feasible (they are referred to as *sterically clashing*), and PLUG can remove them.

It is useful to consider not only voxels in the conformation space of the entire protein, but also voxels in the conformational space *C*(*P*) of a subset *P* of the protein’s residues. Such a voxel is defined only by amino-acid type assignments to the residues in *P*, along with bounds on a subset of the internal coordinates: for example, the conformation of the residues in *P* will not depend on the value of the sidechain dihedrals of a residue *r* ∉ *P*.

If |*P*| = 1 then the voxel is called a *residue conformation* or *RC*^22^. The conformational space of each residue is small enough to enumerate explicitly (often <100 RCs). For example, Ref. 32 tabulates the values of the sidechain dihedral angles for each amino-acid type in a large number of experimentally determined protein structures, and the vast majority of them fall in a small number of voxels in sidechain dihedral space. For purposes of this paper, each cluster in sidechain dihedral space (i.e., each *rotamer*) identified in the Penultimate rotamer library (Ref. 32) is treated as a voxel, and each dihedral is allowed to deviate by up to 9° from the modal value in the cluster.

For |*P*| > 1, *C*(*P*) consists of all possible combinations *v* = *v*_1_∩*v*_2_∩… where *v*_1_, *v*_2_, etc. are RC assignments for each of the residues in *P*. The voxels *v* in *C*(*P*) are intersections of RCs because they consist of the set of conformations bounded by the inequalities defining each of their constituent RCs *v*_1_, *v*_2_,… Thus the number of voxels in *C*(*P*) scales exponentially with respect to |*P*|. As a result, efficient combinatorial search algorithms are necessary to search the enormous sequence and conformational space of the entire polymer, even if conformational flexibility is not modeled for residues that are far from any mutable residues. PLUG is a preprocessing step for this combinatorial search.

### Linear model of steric clashes

Computational prediction of favorable conformations in proteins is still very difficult^33^. However, crystallographic data as well as high-level quantum chemistry calculations have established limits on how close nonbonded pairs of atoms can be found in a biophysical system (except as minuscule contributors to a conformational ensemble). This work will use the lower bounds from Ref. 49 on the distance between two nonbonded atoms of particular element types. These lower bounds include exceptions for hydrogen bonds and for atoms within a few bonds of each other, which are also used here. If continuous flexibility is neglected, it is easy to remove a voxel *v* whose conformations all violate these lower bounds, because *v* consists of a single conformation and inter-atom distances can be computed directly. In the presence of continuous flexibility, it becomes a more difficult problem. Let x = {*x_i_* | *i* ∊ {1,…, *m*}} be the set of internal coordinates bounded in *v*. Thus we can say x ∊ *v* if the vector x of internal coordinate values respects the box constraints that define *v*. Let {*a_j_*} be the set of atoms whose conformation is described by *v*; for example if |*v*| is an RC (single-residue voxel) then this set will only include atoms in that residue. Let *d_jk_*(x) denote the distance between atoms *a_j_* and *a_k_*, as a function of the continuous internal coordinates x. Finally, let *b_jk_* denote the lower bound based on Ref. 49 on that distance; for example if *a_j_* is a carbon and *a_k_* is a hydrogen not within a few bonds of *a_j_*, then *b_jk_* will be the minimum sterically feasible distance between a nonbonded carbon and hydrogen. PLUG aims to determine if there exists a conformation in *v* without steric clashes: that is, x ∊ *v* such that

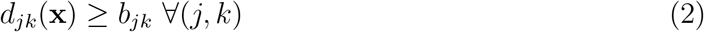

In general this problem is NP-hard even if all the atomic coordinates vary linearly with respect to all the internal coordinates. However, within the constraints of a voxel, we can use an approximation that makes the problem tractable: we assume that interatomic *distances* vary linearly with respect to continuous internal coordinates. For example, within the voxel representing a single rotamer, a 1° increase in the value of a particular sidechain dihedral has roughly the same effect on interatomic distances no matter what the other dihedrals’ values are. The error in this approximation is similar to the uncertainty in the lower bound on interatomic distances (a few tenths of an angstrom: lower bounds on interatomic distances can reasonably be taken as 0.25 to 0.4 angstroms of overlap between van der Waals spheres^49^; see Fig. 1). Moreover, the approximation is often empirically very accurate (see Fig. 2). Areas of conformational space erroneously excluded by the linearized constraint are likely to be missed by an energy minimizer anyway. This is because a convex obstacle like a clashing atom is likely to block the minimizer from exploring conformational space on the other side (see Fig. 2, middle bottom).

**Figure 1:**
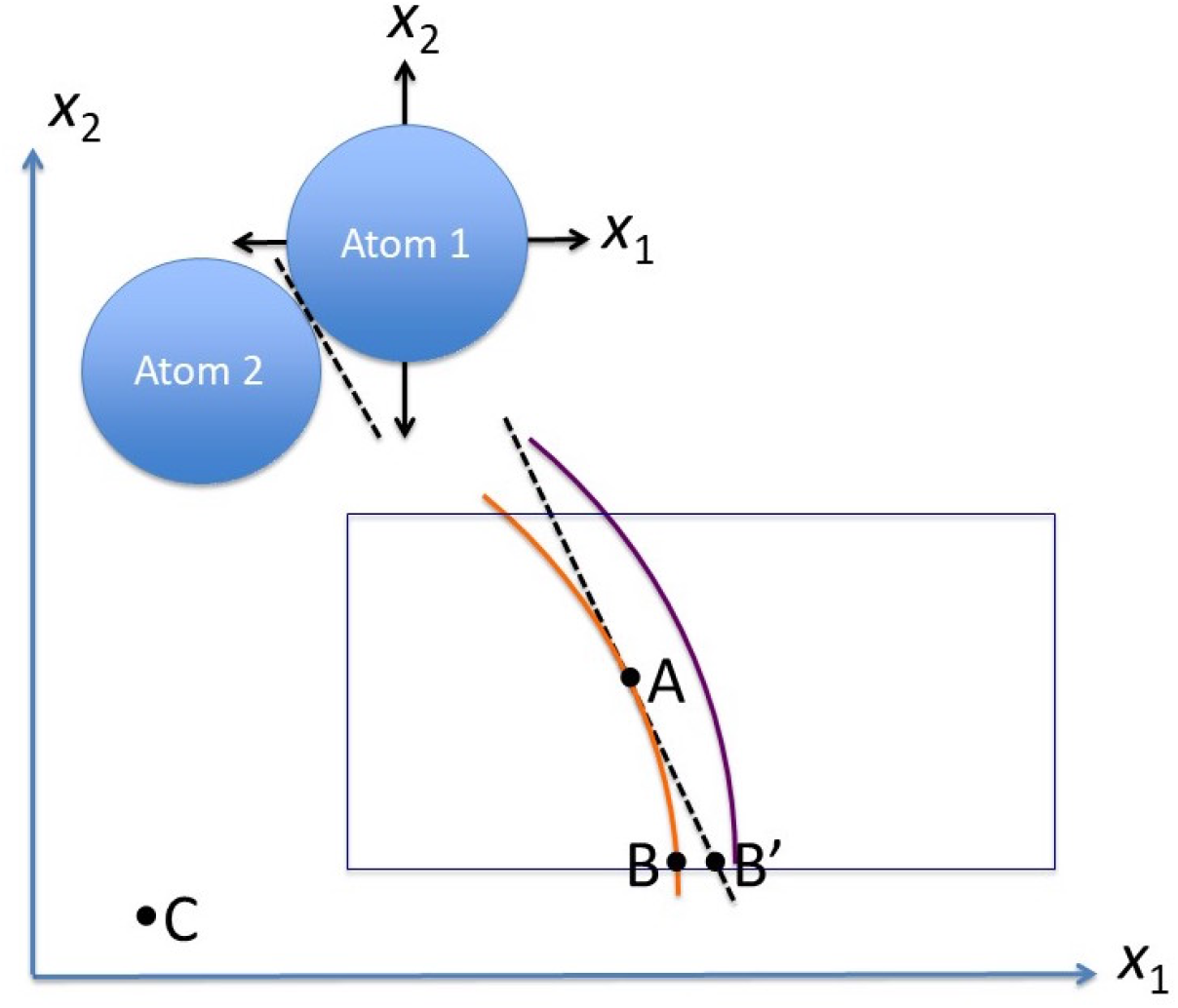
Example of a linearized steric clash, with error estimation. Suppose the position of atom 1 is determined by two orthogonal, continuous degrees of freedom *x*_1_ and *x*_2_. Atom 1 must avoid a clash with atom 2, whose center (in the plane of *x*_1_ and *x*_2_) is at C. The orange arc, centered at C, denotes the boundary of clashing conformational space (by a strict definition^49^), at which atom 1’s van der Waal sphere overlaps atom 2’s by 0.4 Å. The black dotted line is tangent to the orange arc at point A, and is a linear approximation to the orange arc and thus to the clash. The error in this linear approximation within the voxel ranges from 0 (at point A) to the distance between points B and B^′^, at the edge of the voxel. The radius of the orange arc is typically ∼ 1 Å, while point B will generally be ≲0.5 Å from A even with continuous backbone flexibility (see Ref. 19), meaning there are ≲ 0.5 radians of arc between A and B. We can thus estimate the error in the linear approximation as 1/cos(0.5)-1 = 0.14 Å(occurring when B, B^′^ and C are collinear with A and B 0.5 radians apart on an orange arc of radius 1 Å), meaning B^′^ lies within the purple arc, representing a less strict definition of a clash (0.25 Å of steric overlap^49^). Thus the linear approximation itself is a biophysically valid definition of a clash, when used within the bounds of the voxel.

**Figure 2:**
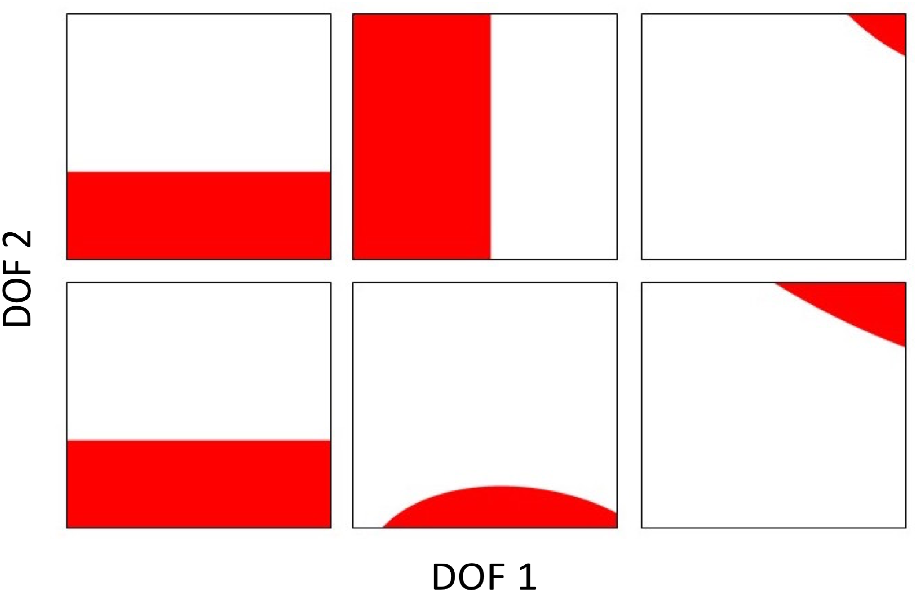
Steric clashes can be approximated as linear constraints. Each graph shows the region of conformational space where a particular atom pair is clashing (red), shown in the plane of the first two continuous degrees of freedom. The length and width of the graph are set to the voxel size. These are the first six atom pairs in the PLUG matrix for the optimal voxel of a design on PDB id 1CC8^42^ from Ref. 20.

Linearized inequalities representing clashes are computed by starting at the center of a voxel (where the minimizer also starts), identifying a point x where *d_jk_*(x) = *b_jk_* by Newton’s method, and then taking the numerical derivative of *d_jk_*(x) with respect to *x* (Fig. 3), yielding a constraint

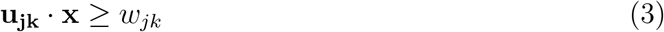

that linearly approximates the true constraint *d_jk_*(x) ≥ *b_jk_*. (The quantities **u_jk_** and *w_jk_* are chosen to provide this linear approximation).

**Figure 3:**
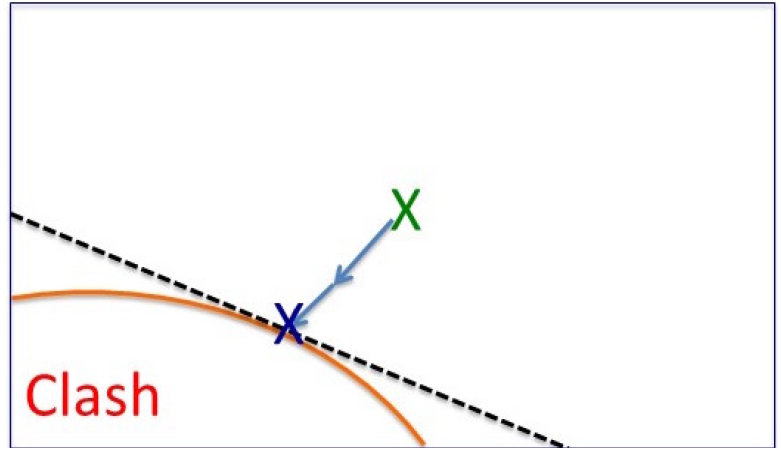
Computation of a linearized clash. Newton’s method starting at the center (green X) of the voxel (dark blue rectangle) leads to a point (blue X) on the boundary (orange) of the sterically clashing region, where a tangent plane (the linearized clash constraint; black) can be computed by numerical differentiation.

Once these linear constraints are computed, the problem of satisfying Eq. (2) becomes a linear programming problem: finding x such that

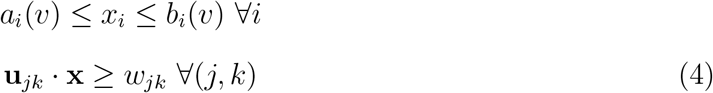

where the first set of constraints just defines the voxel (see Eq. 1). Thus, sterically clashing voxels can be identified by linear programming: these are simply voxels for which the system of inequalities in Eq. (4) is unsatisfiable.

### Matrix precomputations

Since the introduction of DEE^5^ and energy matrices, protein design algorithms have relied on precomputing matrices that tabulate functions of a single RC and/or of a low-order tuple of RCs. PLUG is no exception. First of all, a design with PLUG must begin with the computation of a PLUG matrix: a list of linear constraints representing all pairs of atoms in the system that may clash sterically and give rise to a linear constraint (Eq. 3). Since a pair of atoms can involve at most two different residues, the PLUG matrix can be organized as lists of constraints for each RC and each pair of RCs in the system. Linearized clash constraints for pairs of atoms in different continuously flexible residues are recorded in RC-pair entries of the PLUG matrix; the rest of the constraints (between two atoms in the same residue, or between residues with and without continuous flexibility) are recorded in single-RC entries. Then when PLUG pruning is performed for any (> 2)-residue voxel, the constraints will be looked up for individual RCs and pairs of RCs within the higher-order tuple of RCs defining the higher-order voxel. In other words, if *L* is the list of RCs describing a voxel, then the set of linear constraints to be used for trying to prune the voxel is

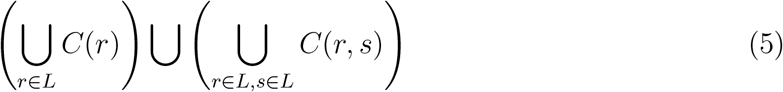

where *C*(*r*) and *C*(*r, s*) are lists of constraints for RC *r* and RC pair (*r, s*) respectively, both of which are looked up in the PLUG matrix. Precomputation of the PLUG matrix is the only part of PLUG that requires explicit modeling of molecular geometries, to identify the coefficients for the linear constraints. Next, pruning of RCs and pairs and triples of RCs can be performed. Like DEE, this results in a pruning matrix, representing what RCs and tuples of RCs are pruned.

### Complexity

Pruning using PLUG is asymptotically more efficient that pruning using DEE. If there are *n* residues with at most *r* RCs each, there are *O*(*n*^2^*r*^2^) pairs and *O*(*n*^3^*r*^3^) triples of RCs to prune, and thus PLUG pruning takes *O*(*n*^3^*r*^3^) time if triples are pruned and *O*(*n*^2^*r*^2^) if not. By contrast, DEE pruning requires comparing a candidate tuple of RCs for pruning to a *competitor* tuple of the same order. So DEE takes *O*(*n*^3^*r*^6^) time with triples: it iterates over *O*(*n*^3^) triples of residue positions, *O*(*r*^3^) candidate triples of RCs given the positions, and *O*(*r*^3^) competitor triples of RCs given the positions and the candidate triple. Similarly DEE pairs pruning takes *O*(*n*^2^*r*^4^) time. The precomputation of the PLUG matrix itself takes *O*(*n*^2^*r*^2^) time, just like computing a pairwise energy matrix, because that is how many pairs of RCs are present in the system. So for both PLUG and DEE it is the pruning of triples (or of pairs if triples are not pruned) that is asymptotically the bottleneck.

### Protocols for using PLUG in protein design

Once the PLUG and pruning matrices have been precomputed, PLUG can be used in a few ways in the protein design computation. First, it can be used as a pre-pruning step just like DEE, removing RCs or tuples of RCs from consideration during combinatorial search. Second, PLUG can be used to prune many-residue voxels later in the design computation. The latter is particularly useful when computing a LUTE matrix^21^, to ensure that the training set for the fitting of LUTE coefficients does not contain inevitably clashing voxels. In this case PLUG pruning is used as part of the pruning check for partial conformations during drawing of sample voxels for the training and validation sets (see Ref. 21). In other words, the LUTE matrix is trained to predict the continuously minimized energies only of non-clashing voxels. This significantly reduces LUTE residuals in the presence of backbone flexibility modeled with CATS^19^ (see Fig. 4). The reason this higher-order pruning is helpful is because a pair of residues may be able to escape a clash when no other residues are around, but the motion that escapes that clash may cause a clash for a different residue pair, once the conformational space of a larger number of residues is taken into account.

**Figure 4:**
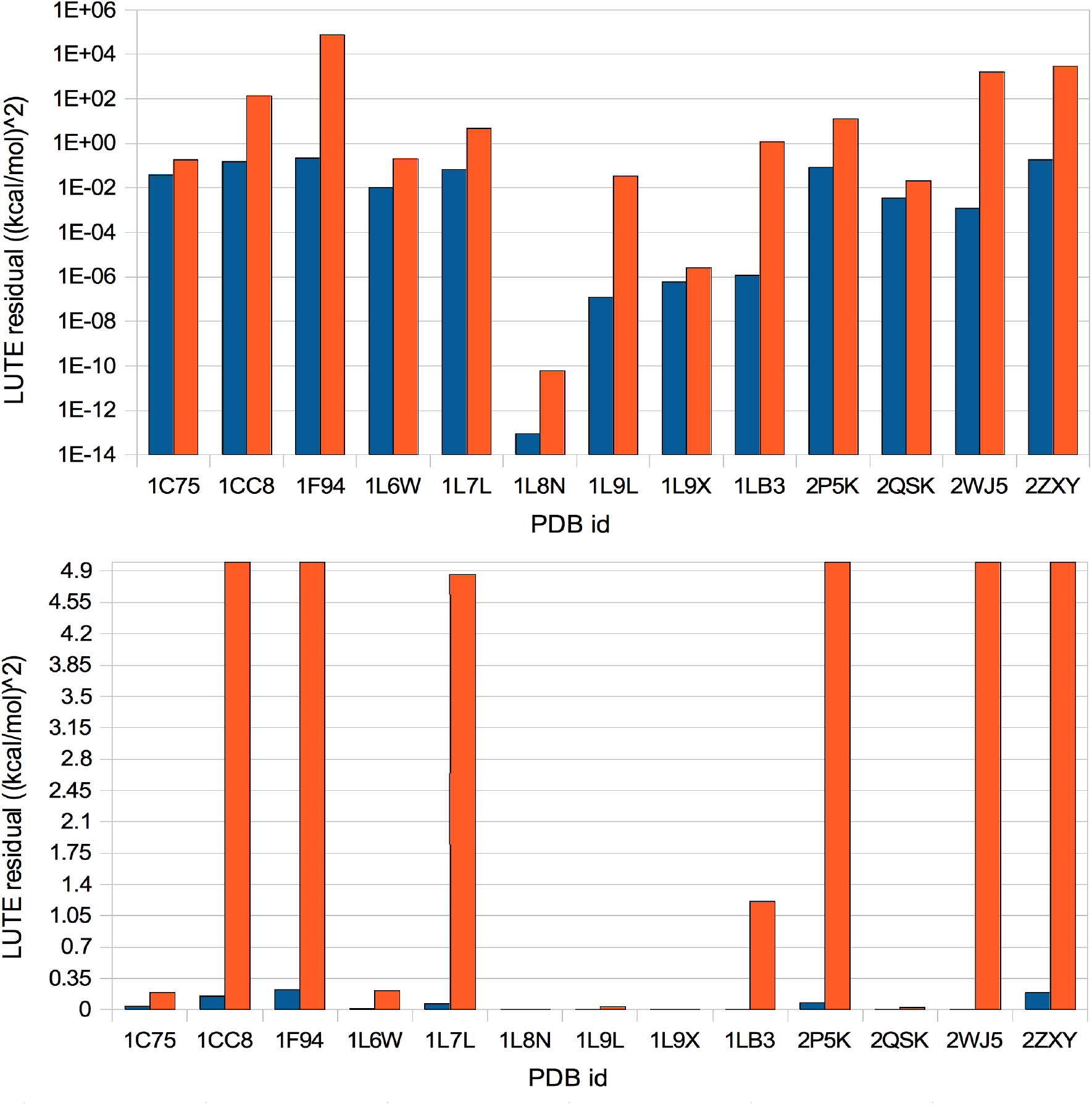
PLUG allows accurate energy fitting using LUTE^21^ for designs with CATS^19^ backbone flexibility. Residuals for LUTE fitting ((kcal/mol)^2^) after pruning with both PLUG and iMinDEE (blue) vs. only with iMinDEE (red); test cases marked by PDB id. Top: logarithmic scale. Bottom: linear scale with horizontal gridline spacing set to the thermal energy at room temperature; thus test cases with residuals below the first gridline (including all PLUG cases) have negligible error, while residuals overshooting this graph likely indicate an unusable fit.

## Results and Discussion

### Pruning power of PLUG

The numbers of RCs pruned by PLUG and iMinDEE were compared at several values of the pruning interval for 96 test cases (Table 1), drawn from Refs. 19 and 20. PLUG pruning is comparable to iMinDEE at a pruning interval of about 5 kcal/mol (Fig. 5), which corresponds to a relatively small design. PLUG is more powerful than iMinDEE at higher pruning intervals, since PLUG pruning is independent of that interval. Up to a pruning interval of ∼15 kcal/mol, iMinDEE and PLUG together prune significantly (∼10–20%) more than PLUG alone. However, designs with many flexible positions (see Refs. 10, 21, etc.) or with continuous backbone flexibility (see Ref. 19) will frequently exceed even a pruning interval of 15 kcal/mol. Moreover, since smaller designs can be accomplished with little pruning, these results indicate PLUG is a suitable replacement for iMinDEE for protein design in general, and can bring the power of iMinDEE to designs in which a pairwise energy matrix and/or pruning interval is not available.

**Table 1:**
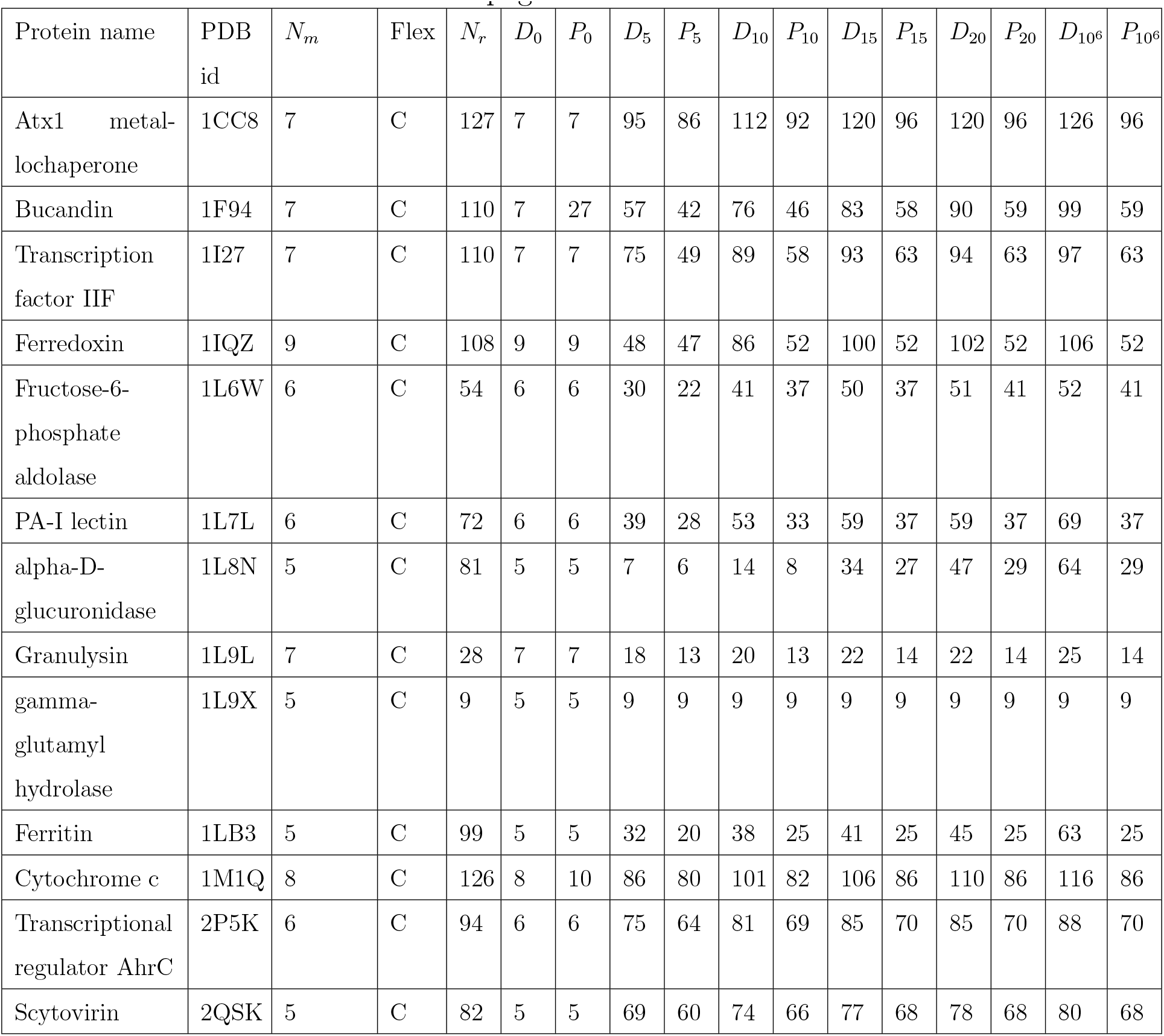
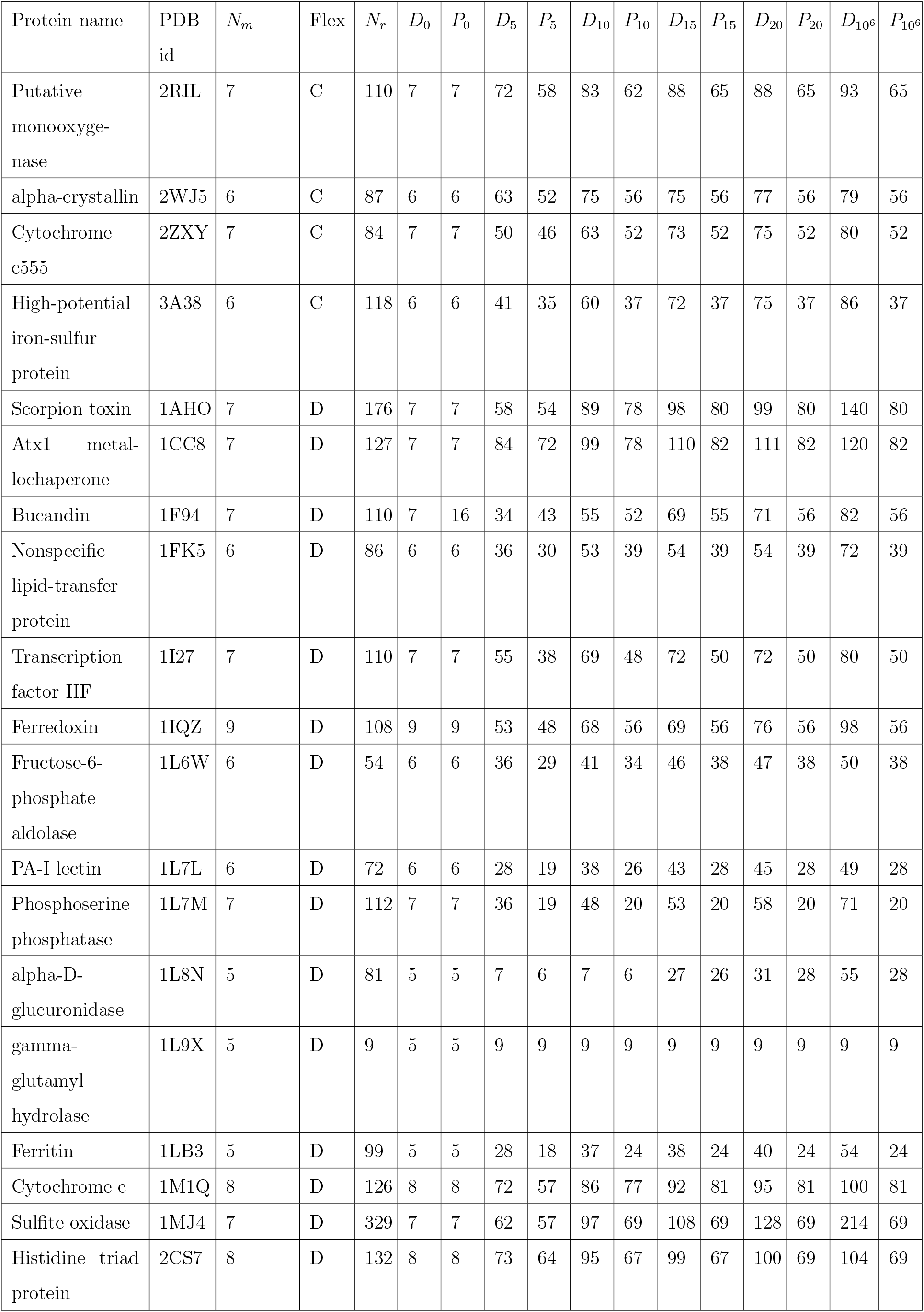
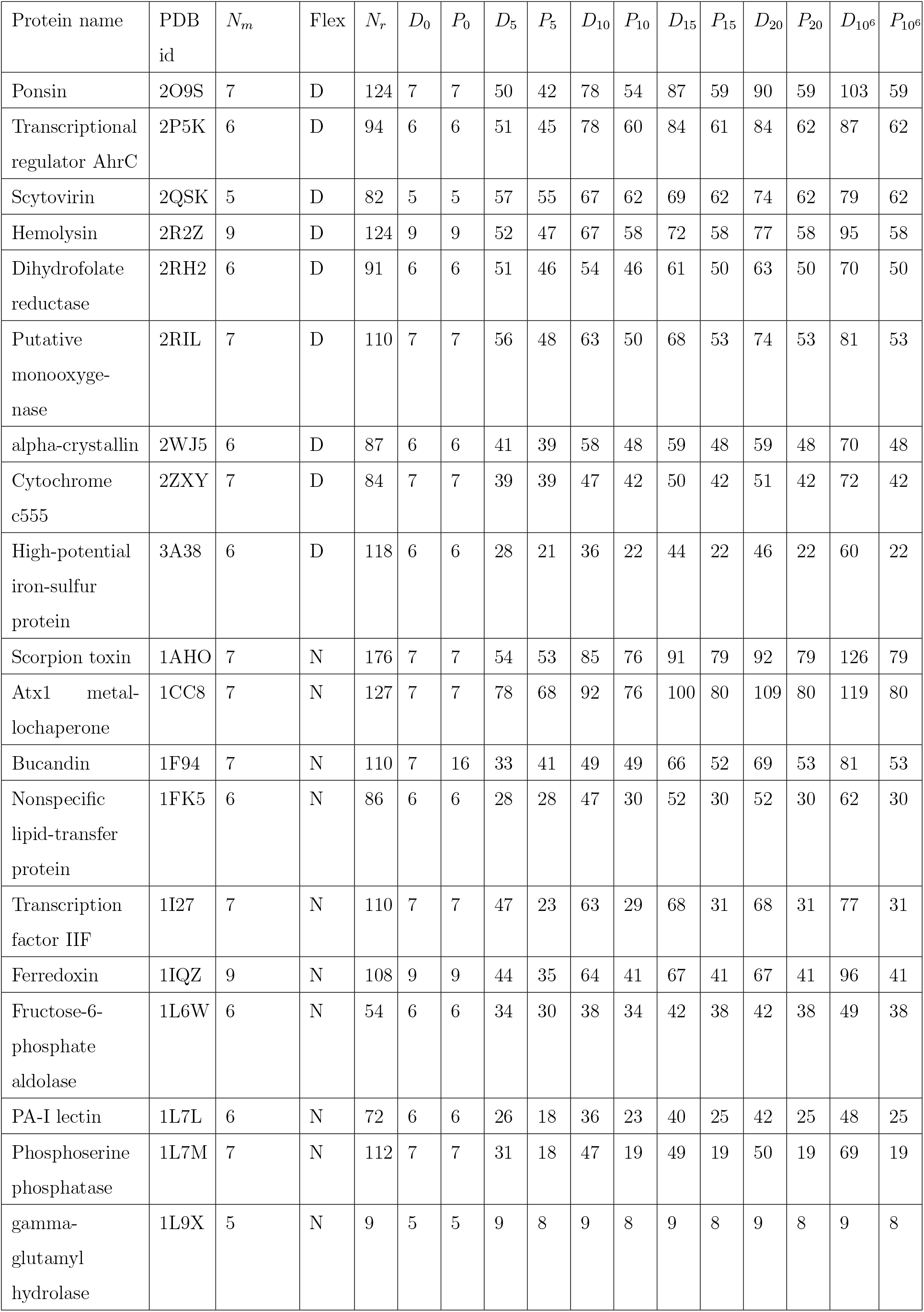
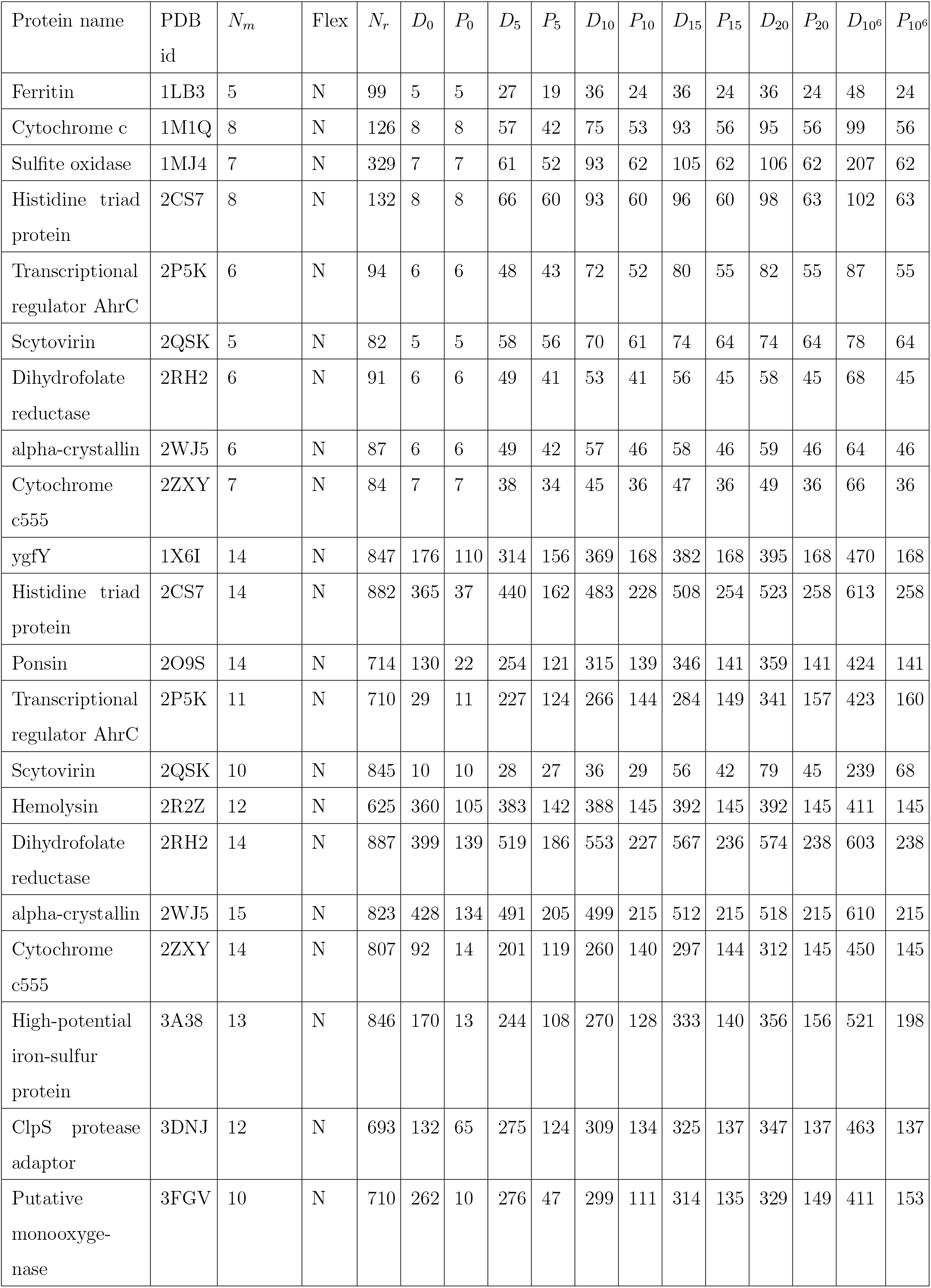
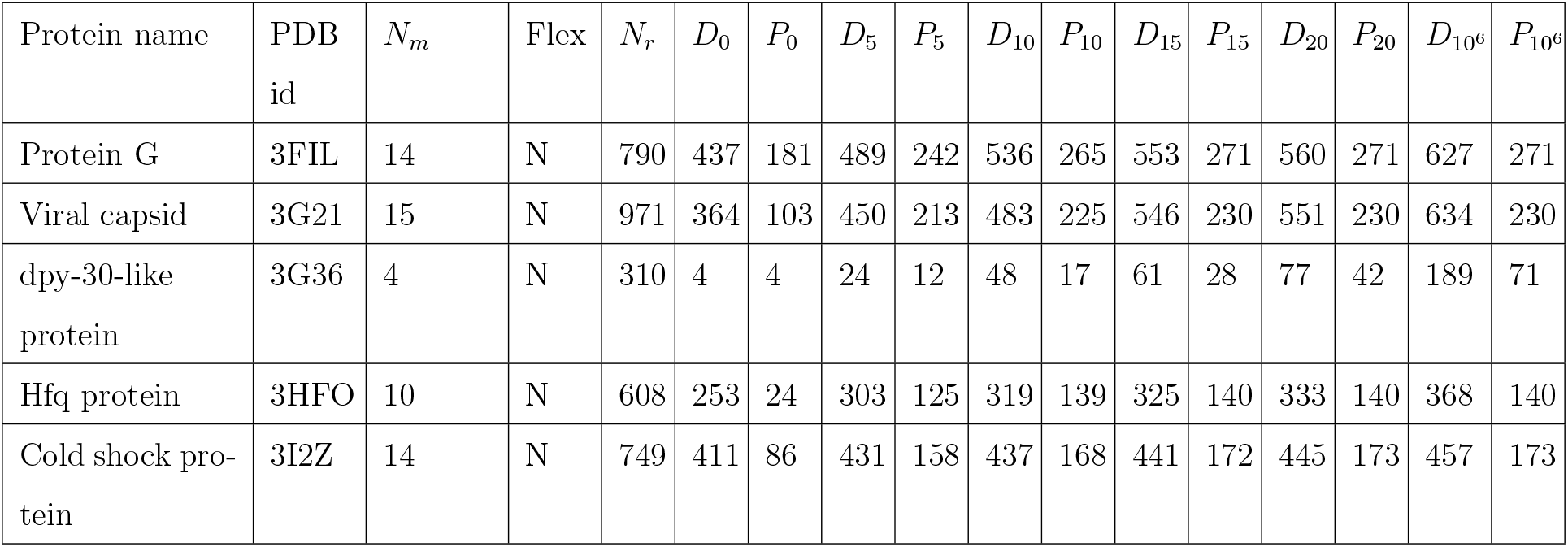
Protein design test cases for examining the pruning power of PLUG and iMinDEE^10^. Each design has *N_m_* mutable residues and a total of *N_r_* RCs across all residues, of which *D_I_* remain unpruned after pruning with iMinDEE using a pruning interval of *I* kcal/mol and *P_i_* remain unpruned after pruning with both PLUG and iMinDEE using a pruning interval of *I* kcal/mol. In every case, *P*_10^6^_ is equal to the number of RCs that PLUG pruning without iMinDEE leaves unpruned, because iMinDEE with a pruning interval of 1000000 prunes only RCs that PLUG alone (which does not use a pruning interval) can prune. “Flex” denotes the level of backbone flexibility for the test case (C=CATS; D=DEEPer; N=None). Test cases where PLUG pruned all conformations are omitted from this table, and instead listed in Table 2. Table continues on next 4 pages.

**Figure 5:**
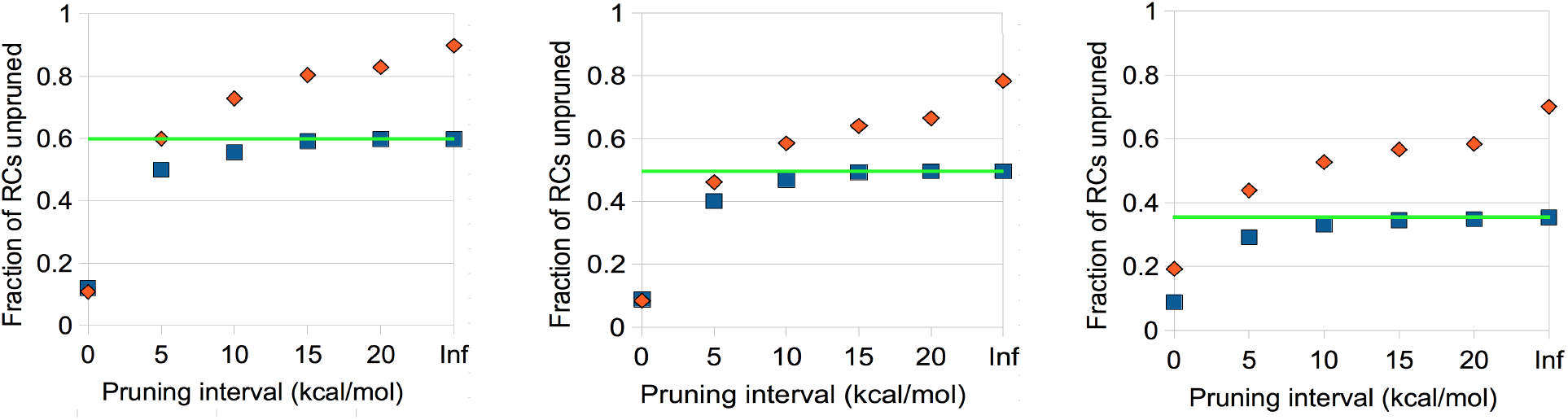
The pruning power of PLUG is similar to that of DEE. Fraction of RCs unpruned after pruning by iMinDEE (red) or iMinDEE and PLUG together (blue), as a function of the pruning interval used for iMinDEE. Pruning was performed using the AMBER energy function^3^, which is pairwise and thus admits iMinDEE pruning (PLUG pruning is not dependent on the energy function). The green line shows the pruning power of PLUG alone, which is independent of the pruning interval (this is equivalent to the rightmost blue dot, because with an infinite pruning interval, iMinDEE only prunes the most obvious steric clashes). Values are the average over 17 design test cases with CATS^19^ backbone flexibility (left), 24 cases with DEEPer^22^ backbone flexibility (middle), or 36 cases without backbone flexibility (right). Test cases where PLUG pruned all RCs are not included.

For 19 of the 96 design test cases tried here (Table 2), PLUG pruned all RCs—it found that steric clashes are unavoidable. Although it is possible to find a minimum-energy sequence and conformation for each of these test cases, this conformation is expected to be biophysically unrealistic (e.g., see Fig. 6). In this sense PLUG improves the realism of protein design. It ensures that the minimum-energy voxel contains a non-clashing conformation. Thus, if PLUG is used in a design, either the predicted minimized-energy conformation will be free of steric clashes, or it will be obtainable by energy minimization starting at a conformation free of steric clashes (and thus any steric clashes will be purely due to the energy function judging other interactions as more important than the steric clash).

**Table 2:**
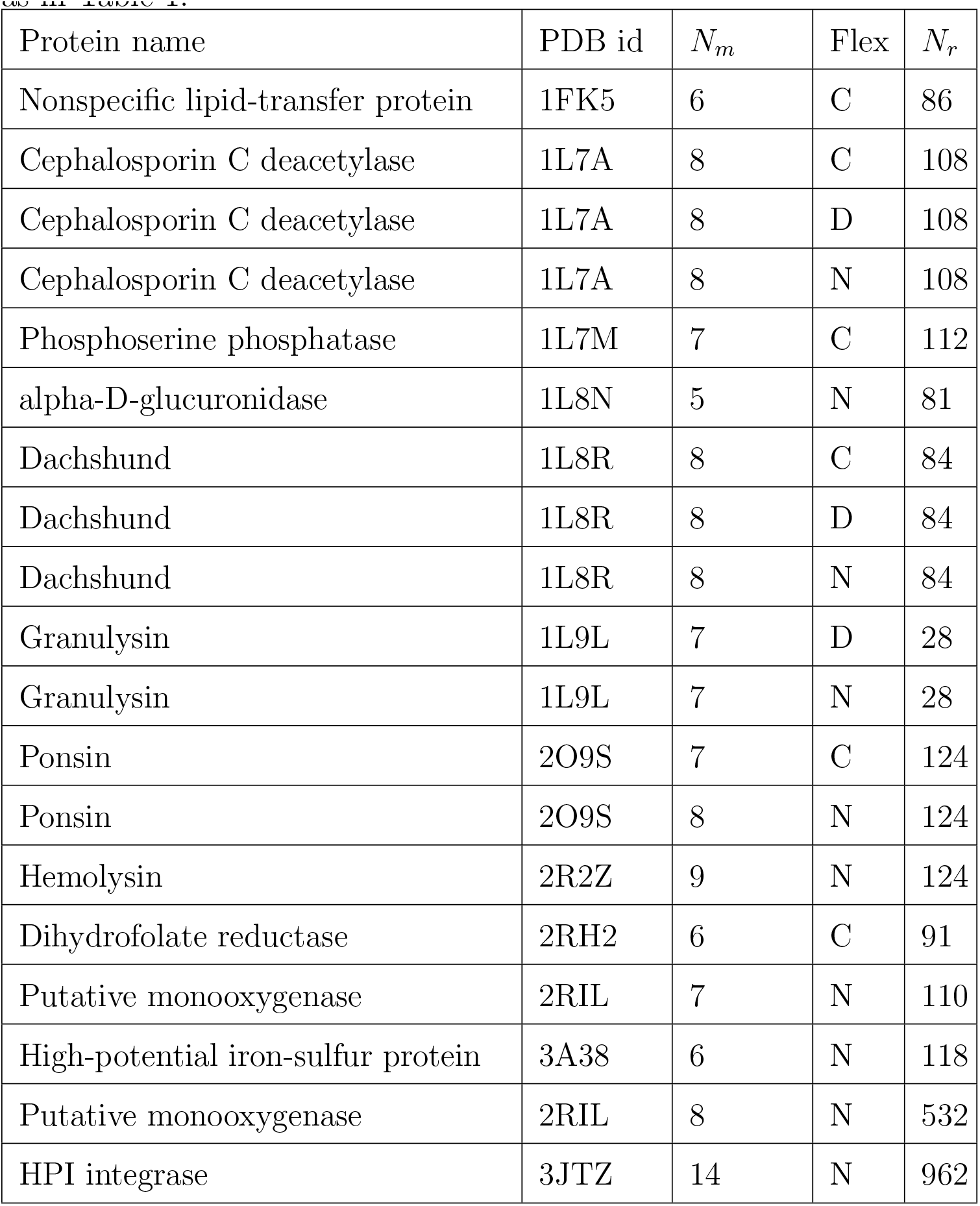
Protein design test cases in which PLUG pruned all conformations. Abbreviations as in Table 1.

**Figure 6:**
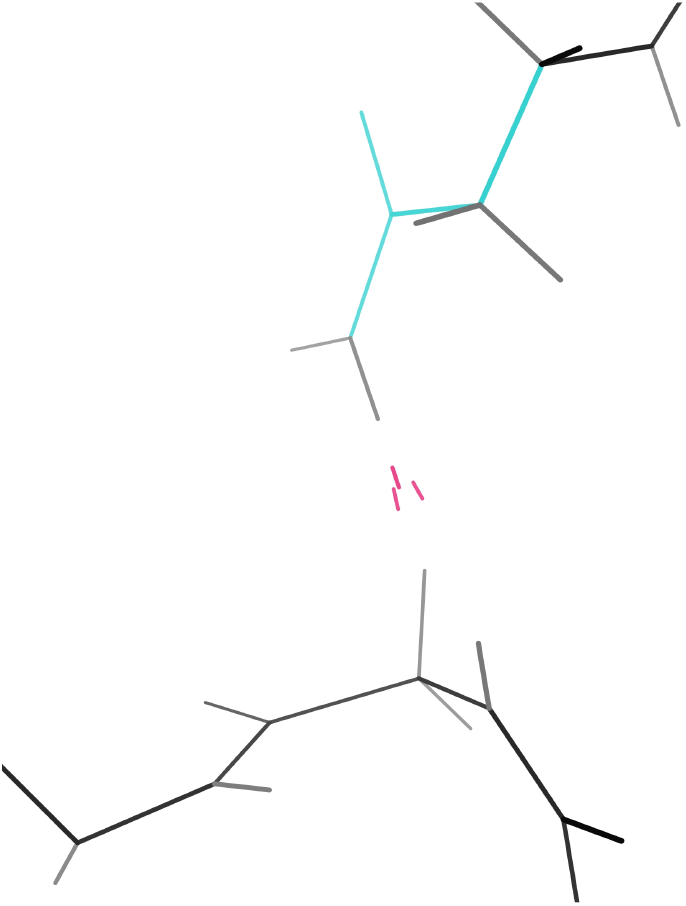
For a design test case deemed impossible by PLUG, the global minimum-energy conformation clashes sterically. PLUG pruned all RCs for the test case from Ref. 19 that redesigned a nonspecific lipid-transfer protein from corn (PDB id 1FK5). When the global minimum-energy conformation was computed without PLUG^19^, a biophysically unrealistic steric clash (pink) is observed.

### PLUG enables the highly efficient LUTE algorithm to incorporate extensive backbone flexibility

The CATS algorithm^19^ greatly increased the level of continuous backbone flexibility available in combinatorial protein design, but at a significant computational cost. Precomputation of a LUTE matrix^21^, which faithfully represents continuously minimized energies despite having the form of a discrete energy matrix, can reduce this cost. Unfortunately, conformational spaces with CATS flexibility have many steric clashes, resulting in high LUTE residuals and impeding the use of LUTE to speed up CATS. Fortunately, PLUG pruning appears to resolve this problem (Fig. 4; Table 3). For each of 13 designs drawn from Ref. 19, a pairwise LUTE matrix matched the CATS-minimized energies of PLUG-allowed voxels to within less than the thermal energy at room temperature (average residual 0.06 (kcal/mol)^2^; maximum residual 0.22). This quality of fit indicates that LUTE can perform the design with negligible error. By contrast, it was impossible to fit a matrix of similar quality to a conformational space that included PLUG-pruned voxels. Using DEE pruning only, only four of the 13 residuals were below the thermal energy at room temperature, and another four were over 100 (kcal/mol)^2^ (i.e., completely unusable). Thus, PLUG is essential for consistent accuracy when performing LUTE computations with CATS backbone flexibility. As a result, PLUG brings the benefits of LUTE to designs with extensive backbone flexibility: efficiency (and thus the ability to handle larger designs), and compatibility with general energy functions.

**Table 3:** Test cases with CATS backbone flexibility for learning a pairwise LUTE energy matrix. The residuals are the mean-square errors of LUTE’s prediction of continuously minimized voxel energies (in kcal/mol). iMinDEE is used in all test cases, while PLUG is used only when indicated.

| Protein name | PDB id | $N_m$ | Residual w/<br>PLUG | Residual w/o<br>PLUG |
| --- | --- | --- | --- | --- |
| Cytochrome c553 | 1C75 | 6 | 0.04 | 0.19 |
| Atx1 metallochaperone | 1CC8 | 7 | 0.15 | 141.95 |
| Bucandin | 1F94 | 7 | 0.22 | 78175.22 |
| Fructose-6-phosphate aldolase | 1L6W | 6 | 0.01 | 0.21 |
| PA-I lectin | 1L7L | 6 | 0.07 | 4.86 |
| alpha-D-glucuronidase | 1L8N | 5 | $8.78 \times 10^{-14}$ | $5.98 \times 10^{-11}$ |
| Granulysin | 1L9L | 7 | $1.23 \times 10^{-7}$ | 0.03 |
| gamma-glutamyl hydrolase | 1L9X | 5 | $6.05 \times 10^{-7}$ | $2.59 \times 10^{-6}$ |
| Ferritin | 1LB3 | 5 | $1.14 \times 10^{-6}$ | 1.21 |
| Transcriptional regulator AhrC | 2P5K | 6 | 0.08 | 12.99 |
| Scytovirin | 2QSK | 5 | 0.0034 | 0.02 |
| alpha-crystallin | 2WJ5 | 6 | 0.0012 | 1680.14 |
| Cytochrome c555 | 2ZXY | 7 | 0.19 | 2979.09 |

### PLUG is necessary for designs where DEE-based pruning is impossible

LUTE can be used to learn a pairwise energy matrix from an energy function that is not pairwise—for example one including Poisson-Boltzmann solvation energies or using the free energy rather than the minimized energy of a voxel^21^. This allows such energy functions to be used in combinatorial protein design calculations, which is otherwise impossible. However, previous applications of LUTE to such energy functions^21^ was limited by the fact that only pruning of the most obvious steric clashes was available. With PLUG, it becomes possible to prune with efficiency similar to DEE for any energy function. Using PLUG as a pre-pruning step, pairwise LUTE matrices were precomputed for six of the CATS designs in Fig. 4 using the Poisson-Boltzmann equation, as implemented in Delphi^34,41^, to calculate the solvation energy for minimized conformations. This energy function is not pairwise, but yielded residuals that were all below the thermal energy at room temperature (Table 4). The average was 0.17 (kcal/mol)^2^, with the highest residual, 0.35 (kcal/mol)^2^, being 0.03 (kcal/mol)^2^ lower than for an equivalent computation with OSPREY’s default, pairwise EEF1^28^ solvation energy function (no DEE pruning, and with CATS backbone flexibility). Without PLUG pruning, LUTE residuals for the Poisson-Boltzmann energy function were unusable (all over 100 (kcal/mol)^2^).

**Table 4:**
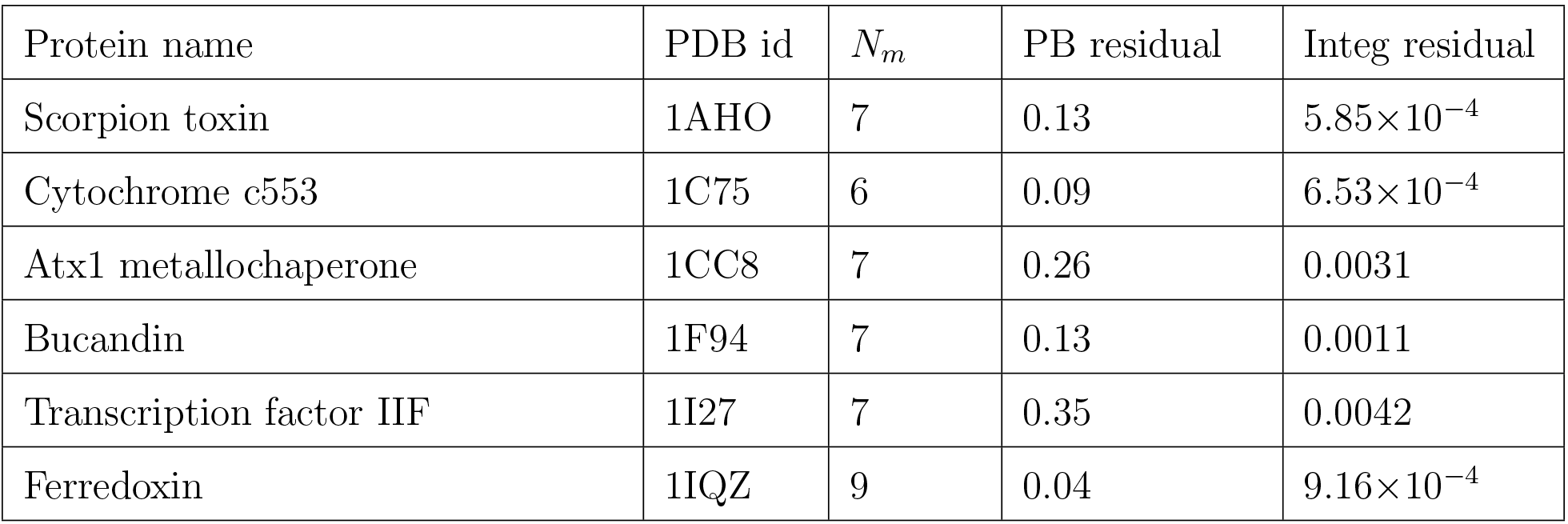
Test cases learning a pairwise LUTE energy matrix using an energy model that does not support DEE pruning–only PLUG is possible here. The residuals are the mean-square errors of LUTE’s prediction of voxel energies (in kcal/mol). For the PB test cases, the voxel energies are the sum of the AMBER energy and a Poisson-Boltzmann implicit solvation term, minimized over voxels using CATS backbone flexibility. For the Integ test cases, the voxel energies are free energies computed by integrating the Boltzmann distribution corresponding to osprey’s default AMBER/EEF1 energy function over a voxel with no backbone flexibility.

A pairwise LUTE matrix was also computed for each of these systems with a rigid backbone but with continuous sidechain entropy. Residuals were all below 0.005 (kcal/mol)^2^ (Table 4). This calculation defines the energy of a voxel differently than usual. Instead of minimizing the energy *E^′^* with respect to the values x of the continuous degrees of freedom within the voxel, the voxel energy is defined as the free energy

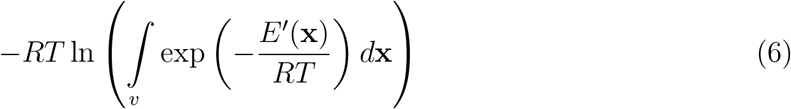

where *R* is the gas constant, *T* is the temperature, and *v* is the voxel. The integral was computed by Monte Carlo integration. Together with the *K^*^* free energy algorithm^13,30^, which approximately discrete partition functions efficiently, this method can be used to compute free energies of binding that include both continuous and discrete entropy.

Also, the procedure for provable computation of a pruning interval^10^, and thus for iMinDEE, is not applicable to multistate designs unless suitable energy constraints are applied manually by the user. iMinDEE for multistate designs is also limited by the fact that it can only use competitors of the same amino-acid type as a candidate RC for pruning (i.e., only type-dependent DEE^51^ can be performed). PLUG suffers no such reduction in power or limitation on provability, and thus will greatly facilitate multistate designs. In this case, the strength of PLUG is that its pruning threshold is based on well-known geometric criteria, not on an energy threshold that must somehow be tuned or known *a priori* to achieve meaningful results.

## Conclusions

The LUTE algorithm^21^ allows efficient incorporation of many types of advanced modeling into protein design. However, it requires a pre-pruning step to remove “ill-behaved” conformations. Previously, this pruning was performed with the DEE algorithm, which imposed significant restrictions on the biophysical model. But now these restrictions are gone: PLUG prunes with similar power to DEE, but with no biophysical assumptions except for well-established knowledge of protein geometry and a validated linearization approximation. The full power of LUTE is thus unleashed. This facilitates great improvements in several areas of biophysical modeling in protein design: energy functions, multistate design, continuous entropy, and backbone flexibility.

## Acknowledgments

I would like to thank Bruce Donald, Jonathan Jou, Marcel Frenkel, and Jeff Martin for helpful discussions, and the TTIC endowment for funding.

